# PUMA: A tool for processing 16S rRNA taxonomy data for analysis and visualization

**DOI:** 10.1101/482380

**Authors:** Keith Mitchell, Christopher Dao, Amanda Freise, Serghei Mangul, Jordan Moberg Parker

## Abstract

Microbial community profiling and functional inference via 16S rRNA analysis is quickly expanding across various areas of microbiology due to improvements to technology. There are numerous platforms for producing 16S rRNA taxonomic data which often vary in file and sequence formatting, creating a common barrier in microbiome studies. Additionally, many of the methods for analyzing and visualizing this sequencing data each require their own specific formatting. As a result, efficient and reproducible comparative analysis of taxonomic data and corresponding metadata in multiple programs remains a challenge in the investigation of microbial communities. PUMA, the Program for Unifying Microbiome Analysis, alleviates this problem in microbiome studies by allowing users to take advantage of numerous 16S rRNA taxonomic identification platforms and analysis tools in an efficient manner. PUMA accepts sequencing results from several taxonomic identification platforms and then automates configuration of data and file types for analysis and visualization via many popular tools. The protocol accomplishes this by producing a variety of properly configured, annotated, and altered files for both analysis and visualization of taxonomic community profiles and inferred functional profiles. PUMA provides an easy and flexible interface to accommodate for a variety of users to produce all files needed for all-inclusive analysis of targeted amplicon sequencing studies. PUMA is an unprecedented open-source solution for unifying multiple microbiome analysis softwares and uses an adaptable implementation with the potential to improve and consolidate the state of microbiome research.

**Body/Findings**

## Introduction

Microbiome research is critical for understanding everything from human health [1] to global ecology [2]. Advances in sequencing technologies and increasing numbers of commercial and academic sequencing providers have made the acquisition of 16S rRNA marker gene microbiome data possible for diverse sets of researchers [3] HYPERLINK “https://paperpile.com/c/FiVobe/3pUu” [4]. Most sequencing providers include basic bioinformatic processing in their pipelines, and provide taxonomic abundance tables and sequence fasta files from the 16S sequences along with the raw data. Unfortunately, many researchers do not have the bioinformatics skills necessary to visualize and further analyze this wealth of data [5]. Packages such as QIIME [6], mothur [7] and phyloseq [8] can be used for analyzing this data, but these require familiarity with coding and the command line in order to use them.

There are several available data analysis and visualization tools that do not require command line, such as the Shiny web app Ranacapa [9] or locally installed programs with graphical user interfaces (GUI) such as STAMP [10] or Cytoscape [11]. These are attractive alternatives for researchers that lack the time to devote to the steep learning curve necessary for installation and use of command line programs [12]. The difficulty here arises in the lack of unity in the data output formats from QIIME or custom commercial and academic pipelines such as MrDNA [13] and Anacapa [14], and the data input formats required for the GUI and web-based analysis and visualization tools. The taxonomy output and metadata formats differ from platform to platform, and may include special file types such as .biom files or delimited text files with specific headings. Reformatting outputs files for each different data analysis and visualization tool is a non-trivial task that may require writing custom scripts or hours of manual work.

Here, we report PUMA (Program for Unifying Microbiome Analyses), a novel tool for comprehensive and efficient streamlining of 16S rRNA microbiome taxonomy data for analysis and visualization. PUMA is a protocol that unifies existing data analysis and visualization tools by formatting common 16S rRNA taxonomic data outputs from a variety of sources to be compatible with the input formats required for multiple basic and advanced microbiome analysis tools. PUMA provides both a Command Line Interface (CLI) as well as a Graphical User Interface (GUI) to accommodate a larger spectrum of potential users. The graphical user interface is ideal for students and or researchers with no experience on UNIX based systems, who are interested in quickly visualizing their 16S rRNA microbiome data. A CLI version is implemented to allow users with UNIX experience, or those who are interested in learning, to customize their analysis and build upon/automate the provided scripts [12]. The graphical user interface is built to be run on QIIME 2 pre-configured virtual machines, upon which a few python custom libraries are automatically installed, to ensure a stable environment for running the program [15].

### PUMA supports input from various microbiome data pipelines

Currently PUMA supports three microbiome raw data processing platforms and/or services: MrDNA, Anacapa, and QIIME 2 [6,13,14]. MrDNA is a commercial full service next generation sequencing provider that offers 16S rRNA amplicon sequencing on a variety of platforms with different price points. Regardless of the sequencing platform MrDNA provides its users with free comprehensive taxonomic analysis in addition to raw data processing using their proprietary pipeline. The pipeline generates operational taxonomic unit (OTU) abundance tables with taxonomic identities and representative sequence files in the .fasta format at each taxonomic level (kingdom, phylum, class, order, family, genus, species).

Anacapa is a software tool kit developed to process environmental DNA (eDNA) sequence data and assign taxonomy data for a variety of marker genes, including 16S rRNA. Anacapa creates a custom reference library for marker genes, generates amplicon sequence variants (ASV), and assigns taxonomies. ASVs have been proposed as a finer resolution replacement for OTU clustering based on sequence similarity [16]. Anacapa provides its users, a detailed taxonomy table with sequences and abundances for each ASV, as well as tables with taxonomies summarized at various percent confidence intervals.

QIIME is a powerful and widely adopted package for processing microbiome data, from raw sequences through taxonomy and data visualization. Tutorials and published protocols are available to walk users through standard data processing [17], but the scope of QIIME may be daunting for novice users, even with the availability of the QIIME 2 Studio graphical interface [18]. It also remains difficult to convert to other analysis/visualization platforms since QIIME provides users with OTU files and sequence files in the .qza format [19].

### PUMA supports inferred functional profile analysis

PUMA also formats taxonomic abundance (OTU or ASV) tables and representative sequence files for prediction of metagenomic content by Piphillin, which uses nearest-neighbor matching of 16S rRNA amplicons and full genomes [20]. Piphillin is currently preferred due to superior performance when benchmarked against other common inferred 16S rRNA functional profiling methods [20,21]. Piphillin has the added benefits of a web interface and of using any standard abundance table and representative sequence fasta file, rather than relying on taxonomic assignments assigned from a specific reference phylogenetic tree, as in PiCRUSt [22]. A drawback to Piphillin is the 10 MB limit placed on uploaded file sizes in the web version. PUMA addresses this by producing subset abundance and fasta files that comply with these limits. The subset files are uploaded to the Piphillin server [23], reference database and percent identity cutoffs are chosen (PUMA currently only supports KEGG [24]), then results are emailed to the user as compressed .tar files. The other drawback to Piphillin is that it provides abundance tables for all predicted genes and pathways (identified by K and KO numbers), but not the associated annotations to assign biological information to the K/KO numbers. The PUMA inferred function protocol also performs queries to the KEGG database in order to properly annotate the genes and pathways returned by Piphillin. Prior to PUMA, this annotation process required command-line experience or labor-intensive manual curation.

### PUMA supports a various types of analysis and visualization platforms

There are a wide variety of research questions that can be addressed using 16S rRNA microbiome data, and the methods used for data analysis and visualization will vary based on the needs of the researcher. PUMA focuses on processing and formatting user data to be compatible with a suite of readily available web-based or graphical user interface (GUI) data analysis and visualization tools. PUMA helps overcome the analysis barriers to entry posed by the requirements of many tools for bioinformatic pre-processing steps or command line coding. Using the PUMA supported tools, researchers can explore data and test hypotheses by linking groups of samples or environmental parameters, otherwise known as metadata, to diversity metrics, community composition, and inferred functional profiles.

To accommodate a broad range of potential research questions, analysis options (from simple to advanced), and visualization types, we have included in PUMA data processing integration with the following suite of tools:

**1. Microsoft Excel** pivot tables are an easy way to begin to summarize the massive amounts of data in taxonomic abundance tables for visualizations of the overall community profile of different samples at different taxonomic levels (phylum, class, order, family, genus, species). It can also be easily used to make simple (non-statistical) comparisons of sample abundances at different taxonomic levels.

- Requires taxonomy table as delimited text file.

**2. Ranacapa** is a user-friendly Shiny web application designed to explore biodiversity using environmental DNA barcode data. It includes interactive visualizations and brief explanations of sequencing depth, alpha and beta diversity, and taxonomy distribution analyses. Ranacapa was developed as an extension of the Anacapa pipeline, but can prove difficult to access from other taxonomic identification platforms, like that of MrDNA [9].

- Requires taxonomy table as *.biom or *.txt. For a *.txt tab-separated taxonomy table, ranacapa requires that a taxonomy column be present with a header named “sum.taxonomy” and that the taxonomy hierarchy associated with each OTU or ASV is in a semicolon separated list.
- Requires metadata as a delimited *.txt file, with the first column of the metadata file containing sample names that match the taxonomy table.

**3. STAMP** (Statistical Analysis of Metagenomic Profiles) is a downloadable graphical interface that can quickly generate publication-quality graphics for differential abundance analysis of either taxonomy or functional pathway data without the need to write code or use command-line interface. STAMP supports parametric and nonparametric statistical hypothesis testing for two-sample, two-group, and multiple-group comparisons. It emphasizes the use of effect size and confidence intervals in assessing biological relevance, and supports a variety of visualizations, including heatmaps, PCA plots, extended error bar plots, box plots, and bar plots [10].

- Requires taxonomy table as tab-separated *.tsv file with strict hierarchical and profile formatting restraints. Many classification tools that produce taxonomies or functional pathways are not strictly hierarchical due to labeling or other errors.
- Requires metadata as tab-separated *.tsv file.

**4. QIIME 2** (Quantitative Insights Into Microbial Ecology) provides numerous interactive and advanced data visualization tools and plugins for evaluation of metagenomic profiles [6]HYPERLINK “https://paperpile.com/c/FiVobe/Mg4G” [18]. Although QIIME can be used for end-to-end data analysis, some researchers may receive data processed by other platforms (e.g. MrDNA or Anacapa) and wish to feed the data back into the QIIME pipeline for analysis.

- Requires a *.qza formatted file for the OTU/ASV file created by importing a *.biom file.
- Requires a *.txt delimited metadata file.
- Requires a *.qza formatted phylogenetic tree to perform phylogenetic beta diversity analysis.

**5. Cytoscape** is a unique open-source locally downloadable tool that enables the visualization of networks between community and functional profiles [11]. Basic network analysis and visualization can be performed with the core distribution, with many additional features available as Cytoscape Apps.

- Requires a *.tsv/*.csv file where the first column is a sample, the second column is the taxonomic/functional identification, and the third column is the taxonomic/functional count or “weight”. The following columns for each row are the metadata features for the sample in the first column.

In addition, PUMA has options to complete data processing such as rarefaction subsampling to normalize for variation in sequence numbers between samples [25,26] using QIIME [6], multiple sequence alignment using MUSCLE [27], and phylogenetic tree construction FastTree [28]. PUMA has the ability to to easily expand to a larger suite of tools as necessary.

### Unifying microbiome data analysis and visualization

Despite, or potentially because of, the increase in the variety and purpose of tools available to analyze 16S microbiome samples, it remains extremely problematic to analyze and visualize data across platforms. Combining different analysis pathways into a single pipeline is a non-trivial task. Each taxonomic assignment platform and analysis or visualization tool may have different data input and output formats that need to be reconciled or significant data pre-processing steps that need to occur before the various analyses can be performed.

For example some sequencing providers, such as MrDNA [13], produce taxonomy abundance tables that must be rearranged in order to be compatible with most visualization programs, but even for those that are in the right general format many tools have specific formatting requirements [13]. For example, the STAMP tool enforces a “strict hierarchy” requirement where no classification of taxonomy can exist at a lower level than one which was left unclassified [10]. The following classification, from phylum to species: “Proteobacteria, Gammaproteobacteria, Enterobacteriales, unclassified, Escherichia, unclassified”, will produce errors in STAMP because the family is unclassified even though the genus is classified. In addition, STAMP requires that all unclassified columns must be labeled so and cannot be left blank [10]. Another tool, Cytoscape [11], requires that each sample identification and taxonomic identification be a unique row where the weight corresponds to the quantity of the given instance in order to create a network type visualization [10]. Web server-based programs such as Piphillin [20] may have file size upload limitations, necessitating sub-setting of the data.

These formatting and processing steps need to be carried out independently on the taxonomy or functional data for each of the desired analysis and visualization platforms (Figure 1A). PUMA provides the solution to these problems by integrating all of the formatting and pre-processing steps required for the platforms and tools discussed in the previous sections into a single unified protocol with an easy installation procedure (Figure 1B). In addition, PUMA is easily expandable by providing the ability to add a new analysis tool or taxonomic ID platform with one added operation.

**Figure 1:**
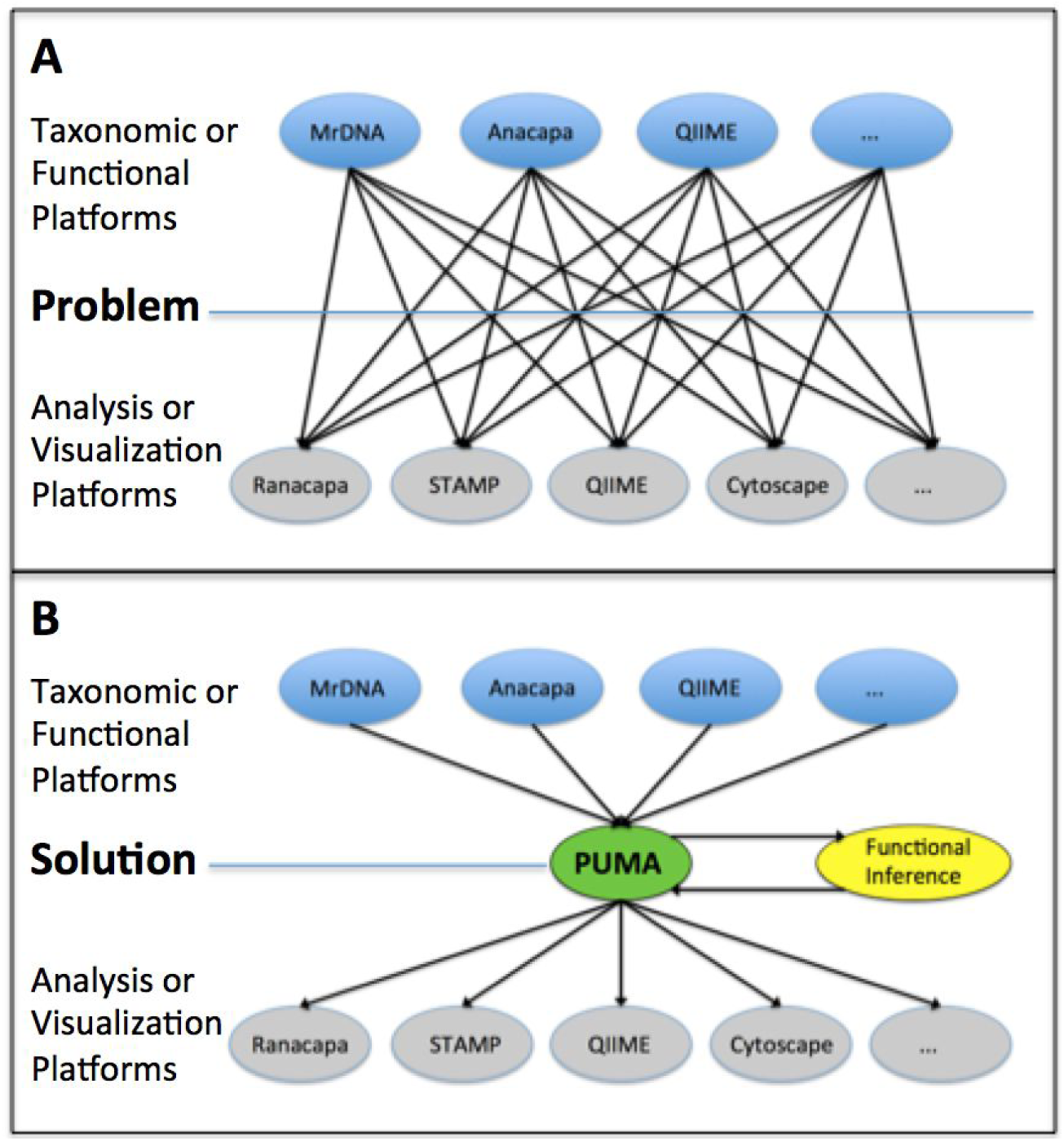
The Problem Presented and the Solution Given PUMA. A) Display of the current problem of lack of unification in the field regarding the different taxonomic ID platforms or functional inference platforms (MrDNA, Anacapa, QIIME, etc.) and their prospective tools (Ranacapa, STAMP, QIIME, Cytoscape, etc.) for analysis producing m x n potential combinations. B) PUMA solves the problem presented in panel A showing how unification can be achieved and scaled simply.

### PUMA use case: undergraduate research curriculum

PUMA was originally developed as part of an upper-division course-based undergraduate research experience (CURE) in microbial ecology at UCLA [29], where it has successfully facilitated students with investigation into a wide variety of research questions. Instructors ran Anacapa 16S rRNA input data through the PUMA program, immediately generating the files necessary for students to use multiple tools for data exploration, analysis, and visualization. This eliminated the need for students or instructional staff to spend hours or days properly formatting files and processing data. Figure 2C illustrates a few representative data analysis visualizations created by students. PUMA has the potential to have a wide impact by making 16S microbiome research accessible for novice researchers, including in educational platforms for research in microbiology [30].

**Figure 2:**
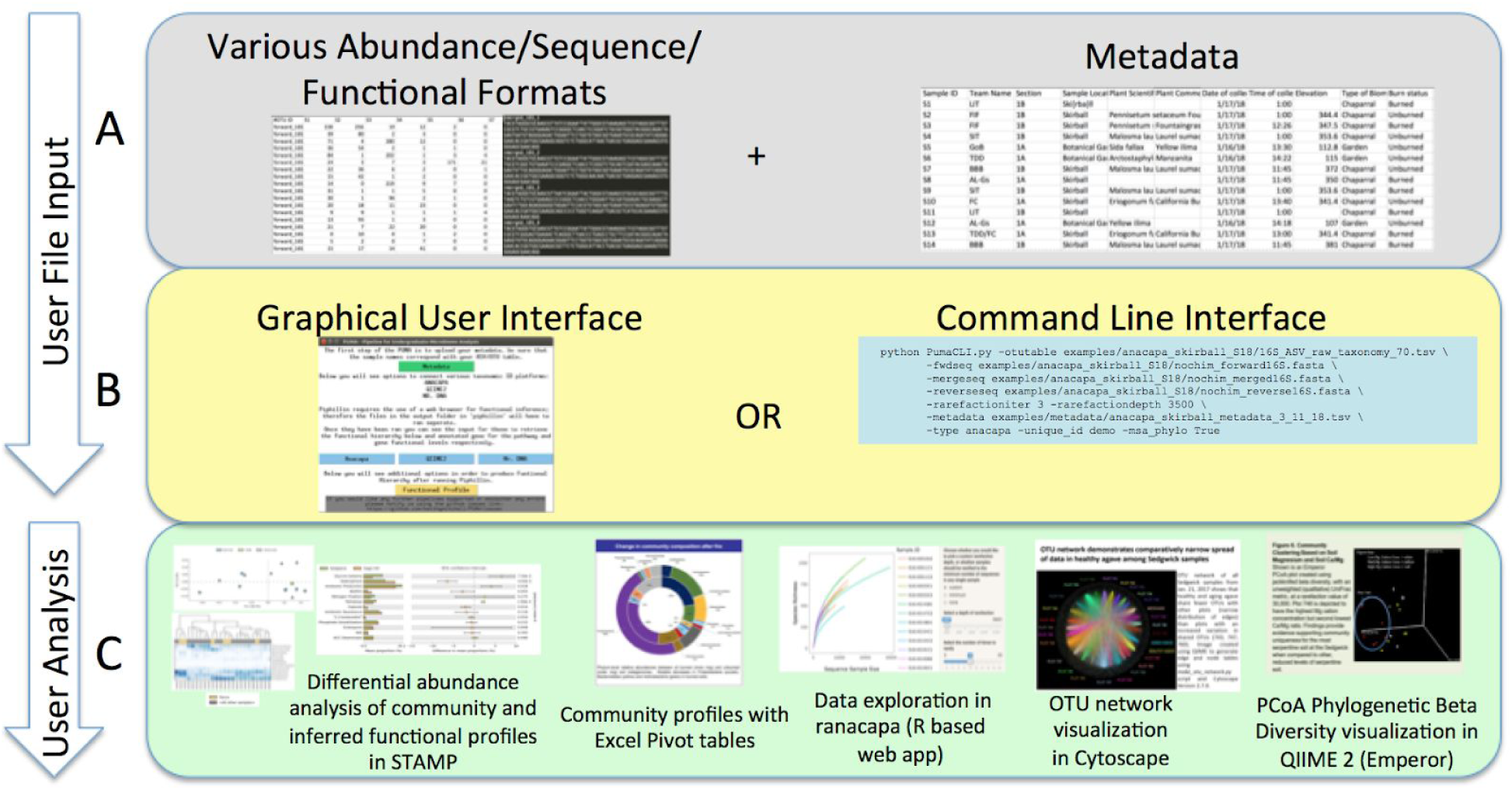
Protocol of the PUMA Software. A) The first panel as part of the “User File Input” displays the simple protocol to be performed by the user such as uploading metadata and various data formats of supported operational taxonomic unit, sequence, and functional file types. B) The second panel as part of the “User File Input” displays the two forms of user interaction with PUMA, through the GUI and CLI, which will enable community and functional profile analysis. C) The last panel as part of the “User Analysis” shows the possible platforms for visualizing community/functional composition data enabled by user input such as STAMP, Excel, QIIME 2, and Cytoscape.

## PUMA Protocol overview

PUMA is distributed as a set of python scripts that is configured to run on a standard QIIME 2 virtual machine (VM) installation. The user executes a single script for both the GUI and CLI versions in order to execute the program. The PUMA protocol consists of two key parts: with part one produces all files for taxonomic community analysis, and part two produces files required for functional analysis. PUMA solves the problem of going from any of the taxonomic identification platforms to the multitude of visualization and analysis tools available, as previously discussed in Figure 1, by enforcing standardized files as part of the unification process (Figure 2).

### Protocol: PUMA verifies metadata

The user uploads their metadata describing the samples, taxonomy abundance (OTU or ASV) table and sequences from any given supported platform. The first part of the PUMA protocol to verifies the metadata and the taxonomy table to be sure the two files have consistent, alphanumeric sample identifiers which are unique compared to other forms of metadata validation [32]. This is a critical step as identifiable metadata is necessary for many downstream analysis steps, and some tools limit the types of characters accepted in the sample identifiers (e.g. underscores, but not periods, are acceptable in sample IDs in ranacapa).

### Protocol: PUMA produces files for community profile analysis

PUMA performs a variety of functions on the taxonomic abundance and sequence files in order to support the suite of tools discussed above. These functions include optional sample rarefaction at a user defined depth and number of iterations (max=10) [33], multiple-sequence-alignment (MSA) by MAFFT [34], phylogenetic tree construction via FastTree 2 [28], and file formatting and annotation for Ranacapa, STAMP, QIIME 2, Piphillin, and Excel. The protocol produces files for community profile analysis in the folder ‘output’, or some other specified directory as an argument in the command line interface. The output folder contains time-stamped subfolders for each PUMA run, each containing subfolders with ready to run files for community profile analyses in Microsoft Excel, STAMP, ranacapa, and Cytoscape. In addition, pre-processed feature table (taxonomy), metadata, and phylogenetic tree files are created that can be imported directly into the QIIME 2 pre-configured virtual machine. A variety of analyses such as alpha- and beta-diversity can be performed in QIIME 2, as well as principal component analysis based on phylogenetic diversity metrics.

### Protocol: PUMA produces files for inferred functional profile analysis

The PUMA protocol consists of three steps necessary for the generation and visualization of inferred functional profiles. The first step is automatically performed at the same time as the generation of the community profile analysis files. PUMA creates a “piphillin” subfolder in the time-stamped output subfolder. This folder contains the original data formatted as a ‘phiphillinotu.csv’ taxonomic abundance table and a ‘phiphillinseqs.fasta’ representative sequence file. If the fasta file exceeds the file size limit of 10 MB enforced by the Piphillin server, PUMA subsamples the data into the number of necessary file sets of ‘.fasta’ and ‘.csv’ files (e.g. piphillinseqs1.fasta; piphillinseqs2.fasta; piphillinotu.csv1.csv; piphillinotu.csv2.csv). Second, each of the sets of Piphillin files in the output directory are uploaded to the Piphillin functional inference web server [23], which returns ‘.tar’ files to the user via email.

Finally, the ‘.tar’ files can then be run directly in the PUMA protocol, which produces files for functional analysis that can be visualized using many of the same tools used for community profile analysis, including STAMP, Excel, and QIIME 2. Importantly, the PUMA protocol also performs queries to the KEGG database using the KEGG genes to pathway API in order to properly annotate the Piphillin gene estimations [35]. The BRITE hierarchy file of the KEGG database is downloaded and used to evaluate the functional hierarchy based on Piphillin pathway estimations. This ensures that estimated gene expression levels and hierarchy levels are inferred using the actively updated information. Annotating the genes and pathway expression from Piphillin is necessary when producing data visualizations with informative identifiers, and greatly reduces the need for manual querying of KEGG.

PUMA produces a timestamped output subfolder for the functional profile files, including a gene description and functional hierarchy file designated for use in STAMP and Excel. This file contains annotated gene names and functional pathways, as opposed to just “K number” identifiers, and vastly increases the efficiency and ease of data analysis and visualization. PUMA also produces weighted functional network files for usage in Cytoscape, which is a platform for visualizing important gene networks between samples.

## Conclusions

PUMA improves the accessibility and comprehensiveness of microbiome investigations by providing users with a simple way to unify the output of various 16S rRNA taxonomic identification platforms with a suite of tools for data analysis and visualization. The protocol accomplishes this by producing properly configured, formatted, and annotated files for analysis of taxonomic community profiles and inferred functional profiles. This process of data manipulation can often be performed by sequencing services for additional fees or completed by users with significant time commitment, both of which could be barriers for those with funding or time constraints. PUMA is an open-source solution which is highly accessible to a wide spectrum of users as it can be used as a graphical user interface (GUI) as well as a command line interface (CLI). The protocol also establishes a standard taxonomic table format with scalable architecture for expanding functionalities to accommodate more taxonomic file formats and programs for visualization and analysis. PUMA provides an easy and flexible interface for a variety of users requiring a clear and brief interface for production of files needed for all-inclusive analysis of targeted amplicon sequencing studies. PUMA has the potential to have a strong, positive impact on research, education, clinical diagnoses, and other fields interested in microbiome analyses.

## Resources

A manual regarding the usage of PUMA for version 1.1 is provided at the following link: <u>PUMA User Manual</u>. The manual is composed of sections which cover installation and running the PUMA graphical user interface and command line interface. The manual will additionally provide resources regarding the usage of tools for visualization supported by the pipeline. The manual additionally has setup instructions for a VirtualBox virtual machine (VM) for ease of user environment replication [36]. Example data for testing PUMA includes example data from the currently supported platforms: Mr DNA, QIIME 2, and Anacapa. The sample data from QIIME 2 is the same as the “moving pictures” human microbiome example dataset available on the QIIME 2 website. The sample data from Anacapa consists of a dataset taken from soil sampled one month after the December 2017 Skirball wildfire in Los Angeles, California.

## Community-oriented software scalability

PUMA is written using a flexible object-oriented system, facilitating the addition of new sequencing services and visualization platforms [31]. The main file ‘master_wrapper.py’ establishes a base class and a class designated by each of the taxonomic identification services can inherit from this base class producing many structural benefits for the software and easier future scalability [31]. The base class contains standard functionalities for processing the standardized metadata, taxonomy, taxonomy counts, and sequences. Each sequencing service has its own class with a unique set of functions that translate the service-provided files into the standard files for the program utilized by the base class. The base class then has a set of functions that it calls to initiate files for the various visualization platforms supported. This structure and organization thus enables a community-oriented system: any user familiar with Python can quickly add their own functionality to the program, from supporting more sequencing service platforms to adding new visualization tools.

## Projected version updates

Support for the Interactive Tree of Life [37], another tool from the makers of iPathway, will soon be provided. In addition, we hope to increase the level of specificity for both the CLI and GUI to accommodate more specifics and customized analysis, such as selecting the visualization platforms desired. The PUMA software may additionally support a web interface for users not interested in CLI features to implement their analysis and visualization. Due to the flexibility of the PUMA interface additions can be easily made to increase the comprehensiveness of the program and accommodate more potential taxonomic identification platforms. For example, other forms of targeted amplicon sequencing may be supported for community profiling along with the obvious potential to scale to more 16S rRNA taxonomic identification and functional inference platforms as well as their respective visualization and/or analysis platforms. Finally, a conda environment will be distributed with the PUMA repository in order to ensure reproducible environment across varying computing platforms [38,39].

## Availability

- Project name: PUMA Version 1.1
- Project home page: <u>PUMA Github</u>
- Operating system(s): Linux (GUI and CLI tested)
- Programming language: Python >= 3.5
- Other requirements: QIIME 2 CLI
- License: FreeBSD
- Manual: <u>PUMA User Manual</u>

